# Single-cell transcriptomics analysis reveals extracellular matrix remodelling in carious human dental pulp

**DOI:** 10.1101/2023.02.15.528696

**Authors:** Anamaria Balic, Dilara Perver, Pierfrancesco Pagella, Hubert Rehrauer, Bernd Stadlinger, Andreas E. Moor, Viola Vogel, Thimios A. Mitsiadis

## Abstract

The carious lesion is a bacteria caused destruction of tooth mineralized matrices marked by concurrent tissue reparative and immune responses in the dental pulp. While major molecular players in tooth pulp decay have been uncovered, a detailed map of the molecular and cellular landscape of the diseased pulp is still missing. Here we used single-cell RNA sequencing analysis, to generate a comprehensive single-cell atlas of the carious human dental pulp tissue. Our data demonstrated modifications in various cell clusters of the carious pulp, such as immune cells, mesenchymal stem cells (MSC) and fibroblasts, when compared to the healthy dental pulp. These changes include upregulation of genes encoding extracellular matrix (ECM) components and the enrichment of the fibroblast cluster with myofibroblasts. Assessment of the Fibronectin fibres’ mechanical strain showed a significant tension reduction in the carious human pulp, compared to the healthy one. Collectively, the present data demonstrate molecular, cellular and biomechanical alterations in the carious pulp tissue, indicative of extensive ECM remodelling and reminiscent of fibrosis observed in other organs.

## Introduction

Carious lesion is a bacteria caused tooth-specific injury, the outcome of which depends on a fine balance between inflammatory responses and the regenerative capability of the dental pulp tissue (Cooper et al., 2014, Duncan et al., 2019). Although the molecular and cellular modifications in the dental pulp tissue upon carious lesion have been extensively studied over the last decades, a comprehensive analysis of these changes is still missing. Histomorphological analyses and gene expression studies have provided a thorough, but limited information on the effects of carious lesion on human dental pulp composition, molecular profile and cellular structure.

Bacterial invasion leads to destruction of tooth mineralized matrices, enamel and dentin, further potentiated by activation of MMPs (<u>M</u>atrix <u>M</u>etalooproteinases), including MMP-2, MMP-9 and MMP20 released from the newly degraded dentin (Chaussain et al., 2013). Proinflammatory responses are triggered in the dental pulp, including the activation of signalling cascades that involve the NFκB pathway and stimulate the expression of proinflammatory cytokines and chemokines, such as IL-1A, IL-1B, IL-6, IL-8, and TNFα (Duncan and Cooper, 2020, Le Fournis et al., 2020). Consequently, immune cells, including dendritic cells and monocytes, are activated and recruited to the site of injury (Hahn and Liewehr, 2007, Duncan and Cooper, 2020, Duncan et al., 2019). These molecular and cellular cascades that constitute the immune response to the carious lesion have been extensively studied (Duncan and Cooper, 2020, Duncan et al., 2019). Pivotal part of the response to carious lesion resides within the heterogeneous dental pulp, composed of various cell types, that differ in origin, function and degree of cell commitment and differentiation, and are embedded in in a complex network of extracellular matrix (ECM) (Pagella et al., 2021, Nanci, 2016). The most abundant cell type present in the dental pulp are the fibroblasts, which regulate extracellular matrix (ECM) formation and turnover, and display high functional and molecular diversity (Jeanneau et al., 2017, Pagella et al., 2021). Dental pulp also contains mesenchymal stem cells (MSCs) that participate in the regeneration and homeostasis of the dental pulp tissue (Sui et al., 2020, Akiyama et al., 2012). Replenishment of the injured pulp tissue by MSCs mostly depends on released cytokines and chemoattractant molecules that coordinate cell migration through the also affected ECM pulp network (Nakashima et al., 2020, Sui et al., 2020).

Pulp ECM is composed of molecules such as Collagen and Fibronectin, and the overal composition and function in healthy human dental pulp tissues is extensively studied and understood (Nanci, 2016, Veis and Goldberg, 2014). However, the caries-induced effects on the biological and mechanical properties of the dental pulp ECM are poorly understood. Generally, repair of soft tissue injury is associated with ECM reorganisation and transient myofibroblast activity that lead to tissue regeneration and restoration of its functionality (Hinz, 2016, Adler et al., 2020). Myofibroblasts share many molecular and functional similarities with fibroblasts and MSCs, and are essential for the ECM turnover during the early stages of injury (Hinz, 2016). However, the extent of ECM remodelling and the potential role of myofibroblast in the carious pulp tissue are not well understood.

In this study we used single-cell RNA sequencing (scRNAseq) analysis to unravel the cariogenic effects on the molecular and cellular composition of human dental pulp, by comparing the transcriptomes of healthy and carious human dental pulp tissues. Our data show significant changes in the dental pulp stromal cells and illustrate a vast remodelling of the ECM network resembling to pathological organ fibrosis. Furthermore, we show important differences in the mechanical properties of ECM between healthy and carious human teeth. Collectively, our data provide the comprehensive single-cell RNA atlas of human carious teeth, and demonstrate the significance of ECM for tissue repair and regeneration upon injury.

## Results

### Molecular profile of dental pulp exposed to carious lesion

Caries is a bacteria caused disease, characterized by enamel and dentin destruction, and consequently changes in the dental pulp, that could lead to secretion of tertiary dentin (Figure 1A-C). There is a great variability in the course and outcome of carious lesions, and the determining factors are not fully understood. We aimed to characterize the molecular landscape of carious lesion that governs cellular behaviours and lesion progression, by performing the single cell RNA sequencing (scRNAseq) on dental pulp cells obtained from carious teeth (Figure 1D and E). In total, 19,305 dental pulp cells from 5 patient samples were evaluated by unsupervised clustering in Seurat v4 (Hao et al., 2021) and visualized in UMAP (<u>U</u>niform <u>M</u>anifold <u>A</u>pproximation and <u>P</u>rojection) space (Figure 1F). Carious pulps contained 5 major clusters previously reported in healthy dental pulp (Pagella et al., 2021) (Figure 1E and H). These clusters include stromal, immune, endothelial, nerve and epithelial cell (Figure 2B), identified at variable sizes in all patient sample (Figure 1G). Approximately 35% of analysed cells (7836 cells) clustered within stromal cell cluster composed of fibroblast (4673 cells) and mesenchymal stem cell (MSC, 3163 cells) subclusters identified by *Collagen type I (COL1A1), Midkine (MDK*) and *RUNX Family Transcription Factor 2 (RUNX2*) expression and *Frizzled Related Protein (FRZB*) expression, respectively (Figure 1I). Furthermore, *Myosin Heavy Chain 11 (MYH11*) and *α-smooth muscle actin (ACTA2*) expression marked a distinct cell populations within MSC subcluster that was clearly distinguishable from the one marked by *THY1* expression (insets in Figure 1I), as previously reported (Pagella et al., 2021). Nearly 30% of total cells (5777 cells) constituted immune cell cluster identified by *PTPRC (CD45*) expression. This cluster was composed of T and B-lymphocytes, monocytes, macrophages and dendritic cells (Figure 1E and Figure 2). Further cell clusters included endothelial cells (4126 cells, or approximately 21%) identified by *Platelet And Endothelial Cell Adhesion Molecule 1 (PECAM-1), CD34* and *Endoglin (ENG*) expression (Figure 1I), and nerve cells, composed of non-myelinating and myelinating Schwann cells previously identified in the healthy pulp (Pagella et al., 2021).

**Figure 1.** Morphological and molecular analysis of carious lesion. **A-C)** Hematoxylin and Eosin (H&E) staining of a carious tooth. **A)** low magnification image showing carious lesion in the crown. **B, C)** high magnification images of the areas marked in A by a black dotted square 1 (B) and 2 (C). **D)** Schematic of the single cell RNA sequencing (scRNAseq) experimental setup, depicting the isolation and enzymatic processing of the dental pulp to a single cell suspension processed for 10x genomics. **E)** Colour coded visualization of cell clusters in UMAP plots. **F)** number of cells analysed by scRNAseq in each carious pulp sample. **G)** Graphical representation of cell fractions per individual cluster, in each carious pulp sample. **H)** Heatmap showing the expression of known markers for the respective cell types. **I)** Distribution of the expression selected markers in the cells visualized in the UMAP space. Abbreviations: d = dentin, Od = odontoblasts, Pd = predentin, TD = tertiary dentin, BV = blood vessel. Scale bar represents 200μm.

We next compared our carious pulp data set to previously published healthy pulp data (Fig 2A). Quantitative analysis revealed no obvious change in the overall size of each cluster in terms of cell proportion between the two data sets (Figure 2B), which could be attributed to high variability between the samples. Gross examination of differential gene expression showed that among the top 25 upregulated genes were genes involved in immune response, namely activation of B lymphocytes, phagocytosis and antigen recognition (Supplementary Table 1 and Supplementary Data 1). Hence, we first analysed the immune cell cluster.

**Figure 2.**
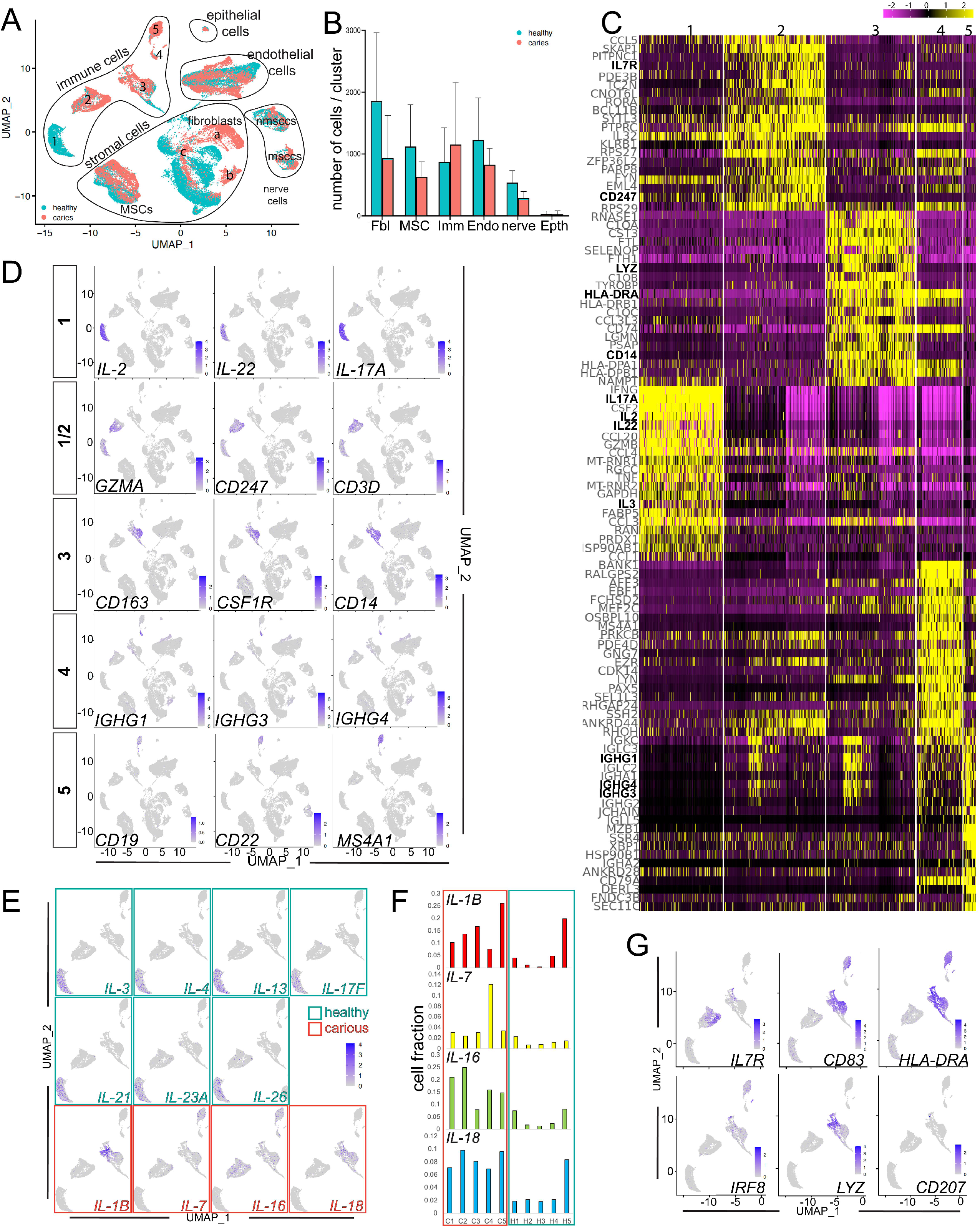
Comparative analysis of the transcriptome profile of carious and healthy pulp. **A)** UMAP presentation of major clusters in carious (red) and healthy (green) dental pulps. **B)** Comparison of cell fractions in major clusters between the healthy (green) and carious (red) pulps. **C)** Heatmap depicting the expression levels of top 20 marker genes for each of the 5 cell subclusters indicated in A. The genes were identified by Seurat. **D)** Feature UMAP plot representation of specific subclusters of immune cells. **E)** UMAP visualisation of the distribution of *IL-3, IL-4, IL13, IL-17F, IL-21, IL-23A, IL-26*, IL-1B, IL-7, IL-16 and *IL-18* in the immune clusters of healthy and carious pulps. Green framed graphs illustrate IL genes expressed mainly in the healthy pulps, and red framed graphs illustrate IL genes expressed mainly in the carious pulps. **F)** Fraction of cells expressing IL-1B, IL-7, IL-16 and IL-18 in the individual samples. **G)** Feature UMAP plot representation of molecular markers shared between the different subclusters of immune cells.

UMAP visualization revealed distinct subclustering of immune cells from carious pulps when compared to healthy pulp cells (Figure 2A). While the immune cells of the healthy pulp were distributed among all the clusters, majority of these cells positioned within the subcluster 1 (Figure 2A, C) marked by high levels of different Interleukins (IL), namely *IL-2, IL-3, IL-4, IL-13, IL-17A, IL-17F, IL-21, IL-23A* and *IL-26* expression (Figure 2A, C, D, E). Quantitative analysis of these markers showed they were mainly absent in the carious pulp data set, that is marked by higher proportion of cells expressing *IL-1B, IL-7, IL-16* and *IL-18* (Figure 2E, Supplementary Figure 2A and B), albeit at variable levels between the carious samples (Figure 2F). Hence, cluster 1 represents mainly healthy pulp immune cells. The transcriptome profile of subcluster 1 shared many similarities with subcluster 2, such as the expression of *Granzyme A (GZMA), CD247* and *CD3D* (Figure 2D), which suggests that subcluster 2 is composed mainly of T-lymphocytes (Karlsson et al., 2021). This is further supported by enrichment of this cluster for cells expressing *IL-7R* that is normally expressed at high levels in T-lymphocytes (Mackall et al., 2011) (Figure 2G). Cluster 3 was marked by the exclusive expression of *CD163, Colony Stimulating Factor 1 Receptor (CSF1R*) and *CD14*, which identify monocyte cell lineage (Karlsson et al., 2021). Other markers, such as *CD83, HLA-DRA*, and *Interferon Regulatory Factor 8 (IRF8*) (Figure 2H), that mark subsets of monocyte, B-lymphocyte and dendritic cell lineages (Karlsson et al., 2021), were also expressed in the subcluster 3. Subcluster 4 was marked by exclusive expression of *IGHG1, IGHG3* and *IGHG4*, that marks memory and naïve B-cells which were detected in insignificant numbers in the healthy pulp data set (Figure 2D and, Supplementary Figure 1C, Supplementary Data 2). Co-localization of *CD83, HLA-DRA*, and *IRF8* (Figure 2H) with *CD19, CD22* and *Membrane Spanning 4-Domains A1 (MS4A1*) (Figure 2D) in the subcluster 5 indicates that this cluster is composed mainly of B-lymphocytes. Collectively, this data demonstrate that immune cell lineages of the healthy pulp are all found in the carious pulp set, but their transcriptome profile is changed. Increased expression levels of specific Interleukin molecules and increased proportion of monocyte/macrophage, dendritic and B-lymphocyte lineages indicate activation of the specific immune and inflammatory responses to bacterial invasion.

Among the top-25 upregulated genes in the carious pulp set was *Serum Amyloid A1 (SAA1*) gene (Supplementary Table 1), an acute-phase protein produced in response to tissue injury, inflammation, and cancer (Malle et al., 2009). This gene was specifically upregulated in the fibroblast population within the stromal cluster (Supplementary Table 1) that was marked by increased proportion of cells expressing genes associated with activated Transforming Growth Factor beta (TGFβ), Tumour Necrosis Factor (TNF) and *Nuclear Factor Kappa B* (NFκB) signalling pathways (Figure 3A). In the fibroblast subcluster number of cells expressing *NFKB Inhibitor Zeta (NFKBIZ), JUND, CCAAT Enhancer Binding Protein Delta (CEBPD), FOSB*, and others was increased more than 4 folds (Figure 3A). Increased number of cells expressing genes targets of activated TGFβ, TNF and NFκB signalling pathways, like *NFKBIZ*, were also observed in the MSC subcluster (Figure 3A). The expression levels of majority of these genes were also increased in the both subclusters of the stromal compartment of carious pulps (Figure 3B and Supplementary Data 3). NFKBIZ (NFKB Inhibitor Zeta) expression is activated by NFKB, establishing a negative feedback loop, as well as IL-1B (Eto et al., 2003). Similar to JUN and FOS genes, it is regulating immune response through interactions with Toll-like receptors (TLR) (Eto et al., 2003, Medzhitov and Horng, 2009). We detected low number of cells expressing individual *Toll-like receptors (TLR*) in the fibroblast cluster of carious pulps. Approximately 5, 7 and 6.5% of cells expressed *TLR1, TLR3* and *TLR4*, respectively (Supplementary Table 2). Albeit low, this number was increased when compared to healthy pulps (Supplementary Table 2). Pulp fibroblasts reportedly also secrete various cytokines, which enables them to further modulate the inflammatory response of dental pulp (Cooper et al., 2014, Coil et al., 2004, D’Souza et al., 1989, Le Fournis et al., 2020). Within fibroblast subcluster, we found approximately 9% of cells expressing *IL-7*, compared to almost 7% in the healthy pulps (Supplementary Figure 2D and Data 2), while only a small fraction of cells (less than 0.3%) expressed *TNFα, IL-6* and *IL-18*, and cells expressing *IL-1* or *IL-8* were not detected (Supplementary Data 2).

**Figure 3.**
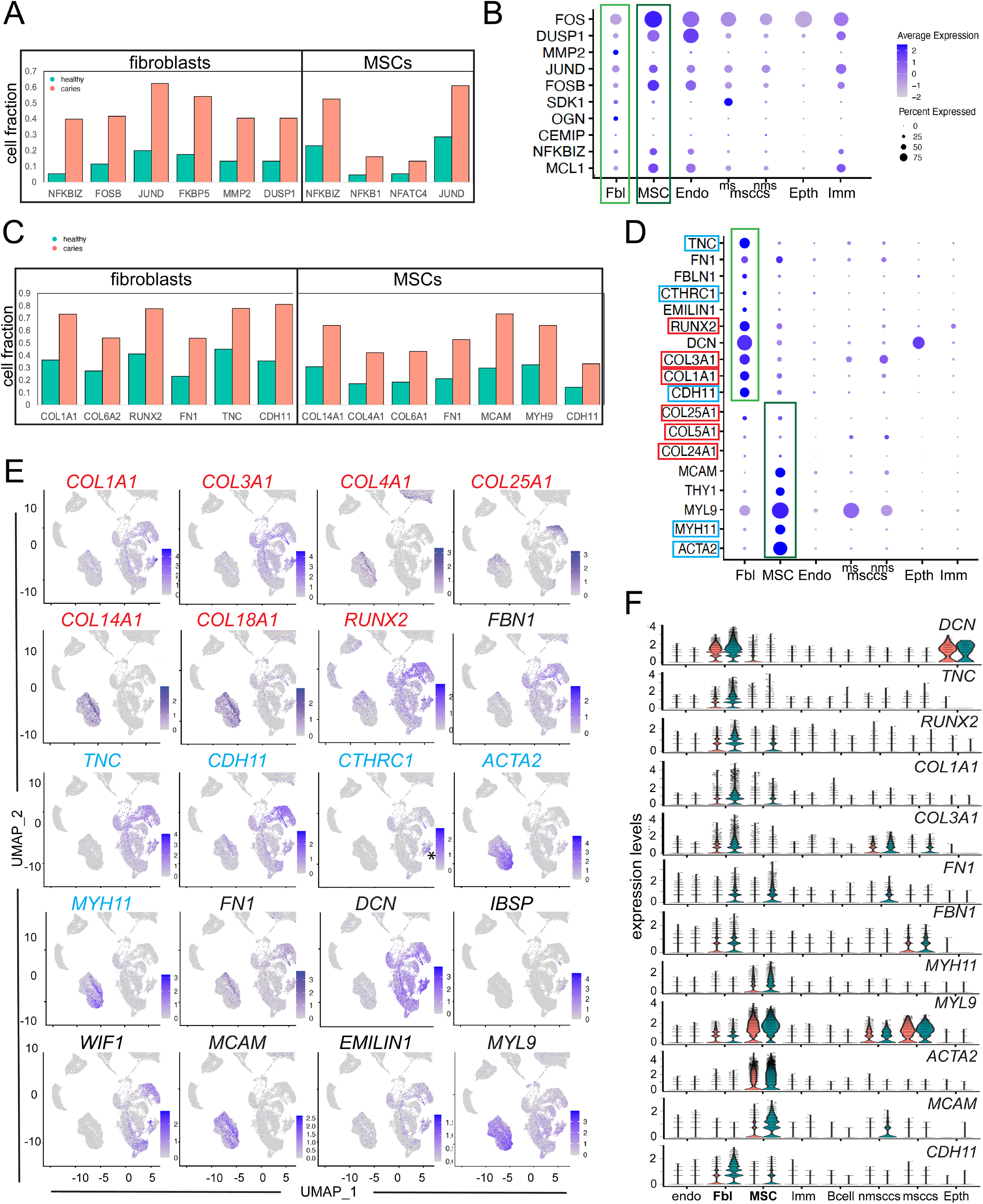
Comparative analysis of the stromal cluster of the carious and healthy dental pulps. **A)** proportion of cells expressing targets of activated NFKB and TGFβ signalling in the specific stromal subclusters of healthy (green) and carious (red) dental pulps, namely fibroblast (left) and mesenchymal stem cell (MSC, right) subclusters. **B)** Dot plot depiction of the expression parameters of selected genes, targets of activated NFKB and TGFβ signalling. **C)** proportion of cells expressing ECM related genes in the specific stromal subclusters of healthy (green) and carious (red) dental pulps, namely fibroblast (left) and MSC (right) subclusters. **D)** Dot plot showing the expression parameters of selected ECM related genes. **E)** Feature UMAP plot representation of the specific cell populations expressing ECM constituent genes (red and black) and myofibroblast-related genes (blue). **F)** Violin plot demonstrating the expression levels of selected ECM-related genes.

More striking change in the fibroblast subcluster was the increase in the proportion of cells expressing various ECM related molecules. There was > 4 fold increase in the number of cells expressing *Osteoglycin (OGN*) (Figure 3B), an ECM associated proteoglycan associated with physiological processes like collagen fibrillogenesis, as well as pathological conditions, such as cancer and ectopic bone formation (Deckx et al., 2016). The expression levels of *OGN* were also significantly increased (logFC 0.68, Supplementary Data 3). We also detected 15% more cells expressing *RUNX2, COL1, COL3A1* (Figure 3C, D, E). In addition, expression levels of these markers increased for over 1.7 folds, compared to healthy pulp (Figure 3F and Supplementary Data 3). Interestingly, the fraction of cells expressing *Integrin Binding Sialoprotein (IBSP*), a marker of osteoblast/odontoblast differentiation (MacDougall et al., 2006), in the fibroblast cluster of carious pulps was approximately 5.4%. This is a 22.5 fold increase when compared to fraction of *IBSP+* cells in the fibroblast cluster from healthy pulps, where only 0.24% expressed this marker (Supplementary Data 2). Collectively, these data suggest the activation of the reparative processes the carious pulps.

Overall, the stromal cluster of carious pulps showed more than 1.5 fold increases in the expression levels, and the number of cells expressing different ECM related molecules, including different Collagen types*, Tenascin-C (TNC), Fibronectin (FN1), Fibrillin 1(FBN1), Elastin Microfibril Interfacer 1 (EMILIN1), Laminin (LAM), Matrix Metalloproteinase 2 (MMP2*), etc. (Figure 3C). Of these, the most interesting was a 2-fold increase in *TNC* expression that was accompanied by an increase from 45% of *TNC+* cells in healthy pulps to approximately 78% of *TNC+* cells in carious pulps (Figure 3C-F). TNC regulates collagen-producing myofibroblasts, which transiently appear in response to injury in various tissues (Hinz, 2016, Imanaka-Yoshida et al., 2020). Myofibroblasts are marked by expression of *ACTA2* and *Cadherin 11 (CDH11*), among others (Hinz et al., 2004, Hinz, 2016). Analysis of the myofibroblast related genes demonstrated distinctly distributed qualitative and quantitative changes between the fibroblast and MSC subclusters. In the fibroblast subcluster, the expression levels and the proportion of cells expressing *ACTA2* was similar between the healthy and carious pulp data sets (Figure 3D-F and Supplementary Data 2 and 3). In contrast, there was over 2-fold increase in *CDH11* expression levels, as well as increase in the number of *CDH11* expressing cells (Figure 3D-F and Supplementary Data 2 and 3). Approximately 22% of fibroblasts from carious pulps expressed another known marker of myofibroblasts, *Collagen Triple Helix Repeat Containing 1 (CTHRC1*) that identifies myofibroblasts in the fibrotic lungs and heart (Valenzi et al., 2019, Ruiz-Villalba et al., 2020). This was a 2-fold increase, compared to healthy pulp, where only 11% of these cells were detected in the fibroblast subcluster (Supplementary Data 2). Interestingly, *CTHRC1+* cells were abundant in the fibroblast subcluster from one carious pulp sample when compared to other samples, both carious and healthy (asterisk in Figure 3E). Recent transcriptomics analyses of the fibrotic skin proposed Secreted Frizzled Related Protein (SFRP) 2 and 4 as a new myofibroblast markers (Tabib et al., 2021). We also observed increase in the proportion of *SFRP2+* cells (~0.9% in carious pulps vs. ~0.3% in healthy pulps) and *SFRP4+* cells (~8% in carious pulps vs. ~4% in healthy pulps) in the fibroblast subcluster of carious pulps (Supplementary Data 2).

MSC subcluster was marked by more than 2-fold increase in the number of cells expressing genes related to ECM organization and remodelling, angiogenesis and vasculature development and wound healing, among others (Figure 3A-D). These include increased fractions of *Melanoma Cell Adhesion Molecule (MCAM), Myosin Heavy Chain 9 (MYH9), FN1* and *COL14A1* expressing cells (Figure 3D-F and Supplementary Data 2), that constituted more than 50% of the MSC subcluster in the carious pulps. In addition, significantly increased *ACTA2* expression marked approximately 75% of cells in carious pulps, compared to ~ 63% in the healthy pulps (Figure 3D-F, Supplementary Data 2 and 3, Supplementary Figure1A and B). Expression levels of other genes associated with myofibroblast phenotype, such as *MYH11 (Ray et al., 2020*) and *Myosin Light Chain 9 (MYL9)* (Layton et al., 2022), were also increased in the MSC subcluster (Figure 3D-F), but the proportion of cells expressing these markers was comparable between the carious and healthy pulp cells (Supplementary Data 2).

It is worth nothing that the changes in the MSC subcluster were accompanied by changes in the fraction of cells expressing *PECAM-1* (0.801 in carious pulps vs. 0.56 in healthy pulps) (Supplementary Figure1A and B, and Supplementary Data 2), as well as *PECAM-1* expression levels (logFC 0.59, Supplementary Data 3). In addition, we also observed increased fraction of cells expressing genes related to cell motility and adhesion (*IGF1R, FBLN1, CXCL12, FN1*, etc.) (Kiely et al., 2005, Kubota et al., 2004, Melchers et al., 1999), and cell apoptosis (*FOXO3, MCL1*, etc.) (Burgering and Medema, 2003, Widden and Placzek, 2021) in both fibroblast and MSC subclusters (Supplementary Data 2 and 3).

### Altered ECM composition upon carious lesion

Transcriptional changes in the expression of ECM protein components suggest potential ECM remodelling and changes in the biomechanical properties of the ECM. Herovici staining of histological sections labels Collagen deposits, and it demonstrated more intense staining in the carious pulps (Figure 4A, B), which was also confirmed by quantification of COL1A1 immunoreactivity that was increased in the carious pulps (Figure 4C-E’). Similarly, immunoreactivity towards other constituents of dental pulp ECM, such as Laminin (Figure 4F-G’), Decorin (Figure 4H-I’), Tenascin C (Figure 4J-K’) and Fibronectin (Figure 4L, M, P) was also increased in carious dental pulp. These data further confirmed our hypothesis of the active ECM remodelling induced by carious lesion. To determine the extent of these changes and their effect on the biomechanical properties of the ECM, we compared the tensional state of Fibronectin in the healthy and carious dental pulps (Figure 4L-Q). Healthy tissues show high tension of Fibronectin fibres, as evidences by a lack of binding of the Fibronectin tension probe FnBPA5 that specifically binds to N-terminal Fibronectin fragments (Fonta et al., 2020). FnBPA5 binding to N-terminal Fibronectin fragments depends on the tensional state of Fibronectin fibres, as their stretching destroys the peptide’s multivalent binding sites (Chabria et al., 2010). In line with this, healthy dental pulp demonstrated no FnBPA5 staining (Figure 4N). In contrast, dental pulp from carious teeth demonstrated relaxed Fibronectin fibres as evidenced by increased FnBPA5 staining (Figure 4O, Q). These data demonstrated changes in the tension of ECM fibres that could affect the reparative processes triggered by carious lesion, as well as the cellular behaviours that determine outcome of carious lesion.

**Figure 4.**
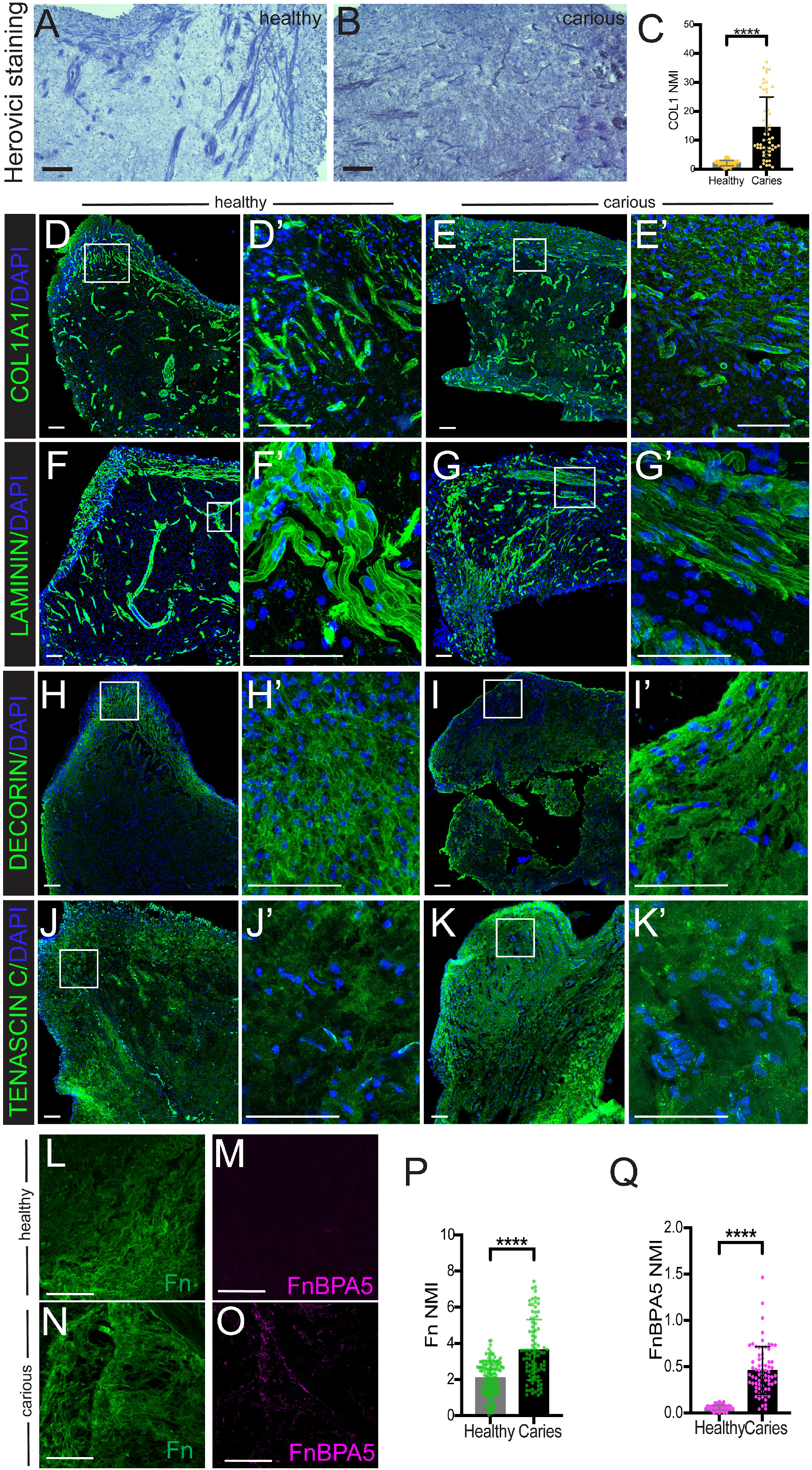
Analysis of extracellular matrix (ECM) changes in the carious pulps. **(A-C)** Herovici histological staining (A, B), and the quantification of COL1A1 fluorescent immunostaining intensity (C) in the healthy (A) and carious (B) pulp cryosections. **(D-K’)** Immunostaining analysis of COL1A1 (D-E’), DECORIN (F-G’), LAMININ (H-I’) and Tenascin C (TNC) protein expression (green) in the cryosections of healthy (D, D’, F, F’, H, H’, J and J’) and carious (E, E’, G, G’, I, I’, K and K’) pulps. DAPI staining (blue) identifies the nuclei. (D’, E’, F’, G’, H’, I’, J’ and K’) are higher magnification images of the areas marked by white square in D, E, F, G, H, I, J and K. **(L-Q)** Analysis of the Fibronectin tension state. Representative confocal microscopy images of cryosections from healthy (L, M) and carious (N, O) pulps immunostained for total Fibronectin (green, L, N) and tension-sensitive bacterial peptide FnBPA5 (magenta, M, O). (P, Q) Quantification of total Fibronectin (P) and FnBPA5 (Q) fluorescent intensity, with data presented as mean + S.D, ****p < 0.0001. Scale bar represents 100μm.

## Discussion

The clinical presentation and outcome of dental caries depends on complex biological, environmental, behavioural and patient specific factors that are poorly understood (Duncan et al., 2019). Various studies have identified individual cellular and molecular changes in carious dental pulp tissues, but the comprehensive analysis of key mechanisms which regulate the course and outcome of this disease are still poorly understood. In this study, we provide a comprehensive map of cellular and molecular changes occurring in carious dental pulps at unprecedented resolution. Our scRNAseq analyses show that carious lesion leads to activation of specific immune responses, generated by activated monocyte/macrophage and dendritic cells and marked by increased levels of cytokines such as IL-1B, IL-7, IL-16 and IL-18, that is accompanied by a unique molecular and structural changes in the stromal pulp compartment marked by upregulation of genes targeted by active NFKB and TGFβ signalling, and a shift in transcriptome signature that resembles molecular changes observed during pro-fibrotic events in various organs. These changes include increases in the expression of myofibroblast-related genes, such as *ACTA2, CDH11, CTHRC1* and others, as well as various ECM protein genes, including *COL1A1, COL3A1, FN1, TNC*, and others, which contribute to ECM remodelling and modifications of mechanical properties.

The first line of defence against carious are odontoblasts that play a role in pathogen recognition, activation of the innate immune response and secretion of reactionary tertiary dentin to resolve the insult (Farges et al., 2015). However, in our data set we have recovered a very small number of odontoblasts, which prevented any meaningful analysis of this cell population. In the remaining pulp tissue we have observed activation of TGFβ, TNF and NFκB signalling pathways, as previously reported (Sloan et al., 2000, Pezelj-Ribaric et al., 2002, Cooper et al., 2014), accompanied by a specific cytokine expression profile. Published RT-qPCR analyses of human pulp tissue exposed to caries report dissimilar cytokine expression profiles, including upregulation of IFN-γ in majority of the tested samples occasionally associated with increased IL-4 and/or IL-10 depending on the depth of carious lesion (Hahn et al., 2000); increased levels of IL-6, L-8, and IL-18, but not IL-1α and IL-1β in carious pulps (Zehnder et al., 2003); upregulated TNF-α, IL-1β, IL-6, and IL-8 (McLachlan et al., 2004) or moderate upregulation of IL-1α, IL-10, IL-13 and IL-17C (Horst et al., 2011). These discrepancies are likely a reflexion of differences in the bacterial pathogen that initiated the caries, diversity of bacterial pathogens associated with secondary bacterial infection and caries progression, as well as the extent/depth and the duration of the carious lesion between different samples (Hahn et al., 2000, McLachlan et al., 2004). Furthermore, these studies indicated complex changes in the cytokine expression profile that affect both pro-inflamatory and anti-inflamatory cytokines. Likewise, our data set revealed distinct cytokine profile, namely increased number of cells expressing IL-1β, IL-7, IL-16 and IL-18, and low to absent numbers of cells expressing *IL-2, IL-3, IL-4, IL-13, IL-17A, IL-17F, IL-21, IL-23A* and *IL-26*. Also, similar to published studies, our data also showed high variability between individual carious and health samples. Majority of these molecules is produced by immune cells, except for the IL-7 that is mainly produced by stromal cells. We have observed moderately increased proportion of cells expressing *IL-7* in the fibroblast cluster, but the most striking increase was presence of *IL-7* expressing cells in the subcluster 5 of the immune cells that was characterized as B-cell subcluster. The available databases (*Human Protein Atlas* proteinatlas.org) also show *IL-7* mRNA expression in the similar population of immune cells.

Whether the observed cytokine profile reflects the response to specific bacterial pathogens remains unanswered, since the microbial and pathogen status of the caries in our samples is unknown. Another possibility is that it reflects the qualitative and quantitative changes of various cell subpopulations within the immune cell cluster. While quantitative analyses of the cellular profile within a sample are limited in scRNAseq data, all the major immune cell populations were present in the carious pulp samples. However, the transcriptome profile of immune cell subsets in the carious pulps was altered from that observed in the healthy pulps, likely reflecting an activation of the immune response to the bacterial infection. Namely, we observed the activation of the dendritic cells and macrophages, that participate in the initial stages of immune response (Hahn and Liewehr, 2007).

While the complex changes observed in the immune cell cluster are interesting, we were more intrigued by the changes observed in the stromal compartment, in which mild carious lesion induced the expression of genes encoding for ECM constituents, namely various Collagens, including COL1 and COL3 that are the main constituents of pulpal ECM, as well as FN1, DCN, TNC and others. Increased number of cells expressing COL1, as well as other genes related to osteoblast/odontoblast differentiation, such as RUNX2 and IBSP, indicates enhanced, ongoing repair in the carious pulp. Interestingly, the observed changes in the ECM composition also share similarities with the fibrotic changes observed in various organs (Weiskirchen et al., 2019). One of the interesting changes was increased number of cells expressing TNC, an ECM glycoprotein involved in physiological and pathological ECM remodelling and repair. Increased TNC expression is observed in inflammation, organ fibrosis (Bhattacharyya et al., 2016, Imanaka-Yoshida et al., 2020, Fu et al., 2017), and in cancer (Midwood et al., 2016). Weak and restricted TNC expression in majority of the healthy adult tissues is transiently induced during wound healing and tissue remodelling, but in pathological conditions, such as fibrosis and cancer, it persists (Bhattacharyya et al., 2022). Among the TNC transcription regulators is TGFβ signalling (Giblin and Midwood, 2015), which is also activated in our carious pulps that demonstrate increased expression levels of TGFβ-1, TGFβ-2 and SMAD3, and higher proportion of cells expressing these genes. Furthermore, studies on heart and skin tissues showed that TNC enhances the myofibroblast phenotype, via TGFβ signalling or through TRL-4, respectively (Katoh et al., 2020, Bhattacharyya et al., 2016). Increased number of cells expressing myofibroblast markers found in our carious pulp samples, is consistent with these findings.

Myofibroblasts share features reminiscent of both smooth muscle cells and fibroblasts, and are important players in fibrotic disorders (Hinz, 2016). While a small proportion of carious pulp myofibroblast-like cells share some molecular similarity with the myofibroblast cells of the fibrotic skin tissue that express *SPRF2* and *SPRF4* (Tabib et al., 2021), our data set indicates more similarity with myofibroblast populations in cardiac and lung fibrotic tissues that is marked by *CDH11* and *ACTA2* expression. Our analyses show that cells expressing these markers constitute a significant proportion in the healthy pulp, as well. This could be attributed to the aging-related changes normally observed in healthy pulps, that include decreased volume and increased fibrous tissue density (Maeda, 2020). Furthermore, our analysis revealed the presence of two distinct myofibroblast-like cell populations in the carious stromal compartment, namely *CDH11* and *CTHRC1* expressing cells in the fibroblast subcluster, and *ACTA2, MYH11*, and *MYL9* expressing cells in the MSC subcluster. Whether there is a developmental hierarchy between the two myofibroblast-like populations remains unclear. The high variability between the different pulp samples hampered further analyses and lineage modelling that could provide a more conclusive answer to this question. The high variability among the samples could be attributed to several factors, including the genetic variability between each individual, and other biological differences, including the age, tooth type, microbial cause, as well as the extent of carious lesion. In this study we aimed to analyse the early carious lesion, which we assessed based on the robust and often unreliable evaluation of the clinical presentation of the caries, namely the extent of the damage in the visible portion of the tooth.

Increased levels of relaxed Fibronectin further indicated enhanced fibrotic changes that accompany pathologically impacted ECM and is found here carious dental pulps (Fonta et al., 2020). Studies in humans and in animal models have shown that Fibronectin plays an essential role during early development, but also in fibrosis (Jarnagin et al., 1994, Muro et al., 2008, van Dijk et al., 2008). Fibronectin regulates different aspects of ECM biogenesis and composition (Sottile et al., 2007), and can contribute to myofibroblast activation (Klingberg et al., 2018). Increased levels of total Fibronectin protein in the carious dental pulps were accompanied by the abundance of relaxed Fibronectin identified by FnBPA5. Increased amounts of relaxed Fibronectin have been previously reported in the cancer stroma in the proximity to myofibroblasts (Fonta et al., 2020). Fibronectin regulates cell adhesion and motility, which determine the speed of wound healing, as well as the invasiveness of cancer cells (Vogel, 2018, Hynes, 2009). Our scRNAseq data show activation of molecular regulators of cell contractility and motility in carious dental pulps, but it is unclear whether they are a direct result of changes in the Fibronectin tension status and ECM remodelling.

The changes in the composition and the biomechanical properties of the pulp ECM induced by carious lesion are similar to those observed in pathological fibrotic tissues of other organs, such as lungs (Valenzi et al., 2019) and heart (Imanaka-Yoshida et al., 2020). One of the surprising findings was the relatively high number of myofibroblast-like cells and TNC expressing cells in healthy pulps. Unlike other soft tissues, dental pulp is contained within the mineralized dentin, which significantly diminishes the stress from physical and mechanical forces of the surrounding tissues (Veis and Goldberg, 2014). One explanation could be that these changes relate to physiological, aging - triggered fibrotic changes that are regularly observed in healthy dental pulp. These changes include diminished cellularity of dental pulp and increased amount of collagen, subsequently leading to remodelling of pulp tissue to adapt to narrowing of the pulp chamber (Murray et al., 2002). Thus, it can be postulated that the onset of aging in the healthy dental pulp might be associated with spontaneous activation of TNC and myofibroblast phenotype. Therefore, dental pulp could be an unique model for studying myofibroblast cells and the onset and molecular regulation of tissue fibrosis under homeostatic conditions, and in the absence of tissue injury or disease.

Collectively, our study provides a remarkable insight into the carious-induced molecular changes in the human dental pulp, that generate a specific cellular and extracellular pulp tissue configuration. Our study clearly indicated the variability between different patient samples that is in line with published data (Veis and Goldberg, 2014) and stresses the necessity for accurate aetiology and classification of caries lesion in order to develop novel, biology-based therapeutic options. Furthermore, similarities between the pulp response to caries and the response of other soft tissues to injury/infection indicate dental pulp as a valuable model to study tissue fibrosis.

## Acknowledgments

We thank Ms. S. Hemmi and Argyro Lamprou for technical help, and Functional Genomics Centre Zürich (ETH / University of Zürich) for the processing and analysis of scRNAseq data. Confocal imaging was performed at the Centre for Microscopy and Image Analysis, University of Zürich, and at the Scientific Centre for Optical and Electron Microscopy (ScopeM), ETH Zürich. This work was financially supported by the University of Zürich.

## Materials and methods

### Human subjects

The procedure for the collection of anonymized human dental pulp and periodontal cells at the Zentrum für Zahnmedizin, Zürich, was approved by the Kantonale Ethikkommission of Zürich (reference number 2012-0588) and the patients gave their written informed consent. Samples were obtained in fully anonymized form from patients of 18-35 years of age. In total 5 tooth extractions were performed by professional dentists at the Oral Surgery department of the Center of Dental Medicine of the University of Zürich. Evaluation of the health status of the tooth was done post-extraction, upon direct observation of the specimen. All procedures were performed in accordance with the current guidelines.

### Cell isolation and single cell transcriptomic analyses

#### Isolation of cells from the dental pulp

Teeth were collected immediately after extraction and placed in sterile 0.9% NaCl, on ice. To obtain dental pulp, each tooth was cracked with a press, and carefully opened with forceps. Isolated dental pulp was placed in a Petri dish filled with cold HBSS and minced into smaller pieces (< 2 mm diameter) for easier dissociation. The dental pulp pieces were transferred to falcon tubes, and dissociated using 5 U/mL Collagenase P (11 213 873 001, Sigma Aldrich, Buchs, Switzerland) in PBS for 40 minutes at 37°C. After incubation in the enzymatic solution, tissue pieces were further mechanically disaggregated by pipetting and centrifuged at 300g at 4°C for 10 min. Cell pellet was resuspended in HBSS containing 0.002% Bovine Serum Albumin (BSA; 0163.2, Roth AG, Arlesheim, Switzerland) and filtered through a 70 μm cell strainer.

#### Single-cell RNA sequencing (scRNA-seq) using 10X Genomics platform

The concentration of cells in the prepared dental pulp single-cell suspensions was adjusted to 1 000 cells/μl. Single-cell library was generated for each dental pulp sample using a droplet method, namely the Chromium System from 10x Genomics, Inc. (Pleasanton, CA, USA), and a Single Cell 3’ v2 or v3 Reagent Kit (10x Genomics, Inc.) according to the manufacturer’s instructions. In each sample, 10 000 cells were loaded into the 10X Chromium controller and the resulting libraries were sequenced in an Illumina NovaSeq sequencer according to 10X Genomics recommendations (paired end reads, R1=26, i7=8, R2=98) to a depth of 50.000 reads per cell.

The gene expression count matrix was computed using Cellranger v7. All data analysis was performed using Seurat v4 (Stuart et al., 2019) and R version 4.2. Clusters were visualized using uniform manifold approximation and projection (UMAP) (McInnes et al., 2018). Samples were initially analysed separately, and data was scaled and transformed using SCTransform (Hafemeister and Satija, 2019) for variance stabilization. The joint analysis of all samples was performed by integrating data using Seurat. All statistical analysis of differential expression was performed on unintegrated data. Differential expression analysis was performed using the Wilcoxon rank sum test. All p values reported were adjusted for multiple comparisons using the Bonferroni correction.

#### GO analysis

To characterize the molecular changes found in scRNAseq dataset, and to put them in a context of biological, cellular and molecular processes, we used g:Profiler, a simple user-friendly web interface with powerful visualisations (Raudvere et al., 2019)

### Tissue processing and analysis

#### Tissue processing and embedding

Healthy and carious human dental pulps were fixed with 4% paraformaldehyde (PFA) at 4°C overnight, immediately following the isolation. The following day, tissue was immersed in 30% Sucrose and kept at 4°C until sinking. Sucrose infused tissue was then transferred to Tissue-Tek O.C.T. Compound (4583, Sakura, Alphen aan den Rijn, Netherlands), snap frozen and serially cryosectioned in 10μm thick sections.

#### Herovici staining

Young and mature COL1 were differentiated by Herovici staining performed on tissue cryosections washed with PBS, following kit instructions (18432, MORPHISTO GmbH, Offenbach am Main, Germany). The stained slides were gradually dehydrated and mounted with Eukitt Quick-hardening mounting medium (03989, Sigma-Aldrich).

#### Immunostainings

Cryosections of human dental pulps were washed with PBS, incubated with blocking solution (1%BSA, 10% donkey serum, 0.5% Triton in PBS) for 45 min at room temperature, followed by incubation with the primary antibody at +4°C, overnight. Primary antibodies used were as follows: ACTA2 (αSMA, ab5694, 1:200), COL1A1 (ab34710, 1:100), COL3A1 (ab6310, 1:200), CD31 (PECAM, ab9498, 1:100), LAMININ (ab11575, 1:100), DECORIN (ab175404, 1:100), TENASCIN C (LS-B8789/74286, 1:50). After incubation with the primary antibody, sections were extensively washed and incubated with appropriate secondary antibody (1:500) for 45min at room temperature. The secondary antibodies used include donkey anti-rabbit (AF488, A32790, Invitrogen) and donkey anti-mouse (AF647, A31571, Invitrogen). After incubation, slides were washed with PBS, mounted using Vectashield with DAPI, and imaged with Leica STELLARIS 5 automated upright confocal laser scanning microscope equipped with HC PL APO CS2 20x/0.75NA Air and HC PL APO CS2 63x/1.3NA Glycerol objectives.

### Analysis of the Fibronectin fibres’ tension state

#### Immunostaining for Fibronectin, FnBPA5 and COL1A1

Tooth pulp cryosections were stained with Fibronectin and FnBPA5 probe as previously described (Fonta et al., 2020). Briefly, thawed sections were washed with ice-cold 1xDPBS and blocked with 0.3M glycine animal-free blocking buffer (Animal-Free Blocker® and Diluent, R.T.U. SP-5035-100) at room temperature for 30 minutes. Sections were then incubated with 1μg/ml Cy5-FnBPA5 or Cy5-labeled scrambled-FnBPA5 (scrambled-FnBPA5) for 1h, followed by three 5 minute washes with Dulbecco’s phosphate-buffered saline (DPBS), second blocking with the blocking buffer and incubation with polyclonal rabbit anti-Fibronectin (ab23750, Abcam) and monoclonal rat anti-Collagen I (7025, Chondrex) antibodies, overnight at 4°C. The following day samples were extensively washed with DPBS before 1h incubation with secondary antibodies and 2 μg/ml DAPI at room temperature. The secondary antibodies used were goat anti-rabbit (AF488, A11034, Invitrogen) and goat anti-rat (AF546, A11081, Invitrogen) antibodies. Following extensive washes after incubation with the secondary antibody, the samples were mounted using DAKO Fluorescence mounting medium (DAKO, Denmark), and imaged.

#### Confocal laser scanning microscopy (CLSM) Imaging

Stained tissue sections were imaged either as a whole section overview (tile-image), or as a randomly chosen high magnification images. Tile-images were acquired as z-stacks using a Leica SP8 confocal microscope with HC PL APO CS2 20x/0.75 Air objective with 0.75 NA, and stitched together using the Mosaic Merge stitching function of LAS X software. DAPI, Fibronectin, Collagen I, and FnBPA5 channels were imaged with 512 x 512-pixel resolution, and the laser power was kept constant. For high magnification images HC PL APO CS2 63x/1.NA Oil objective was used.

#### Image analysis

Quantification analysis of the fluorescent intensity was performed using Fiji software. Control tissues incubated with Cy5-labeled scrambled peptides and secondary antibodies were used to determine the threshold values for each channel. Quantification was performed in randomly selected areas (50 – 100 μm x 50 – 100 μm) of tile-images, and on high-magnification images. In each area/image quantified, the individual signal intensity above the threshold was evaluated in every virtual section (n≥6/z-stack). The FnBPA5-Cy5 and Fn-AF488 intensity ratios were also determined in every virtual section. The collected intensity values for each signal were normalized to DAPI mean intensity. Data were collected as experimental triplicates from 2 healthy and 2 carious pulp tissue samples.

#### Statistical analysis

Statistical analysis was performed in Graph Pad Prism (version 9.4.1) using unpaired student’s T-test with Welch’s correction for p < 0.05.

---

**Supplementary Figure1. Analysis of ACTA2 and PECAM-1 expression in the carious pulps.** Immunostaining of the cryosections of healthy (A) and carious (B) pulps for ACTA2 (green) and PECAM-1 (red) (A, B). Nuclei are marked by DAPI staining (blue). Individual channel images for ACTA2 (a, b), PECAM-1 (aa, bb) and DAPI (aaa, bbb) expression from A and B. Scale bar represents 100μm.

**Supplementary Figure2. Analysis of immune cell related genes. A, B)** Fractions of cell populations in the immune cluster expressing IL genes, in healthy (green) and carious (red) pulps. Cell fractions of specific cell populations found mainly in the immune cluster of healthy dental pulp (A), or expanded in the carious pulps (B). **C)** Cell fractions characterised by the expression of molecular markers also upregulated in the immune cluster of carious pulps. **D)** UMAP visualisation of the IL-7 distribution in the fibroblast subclusters of healthy and carious pulps.

**Supplementary Table 1.** Top25 upregulated genes in the carious pulp data set.

| cluster | gene | p_val | avg_log2FC | pct.1 | pct.2 | p_val_adj |
| --- | --- | --- | --- | --- | --- | --- |
| Immune - subcluster5 | IGKC | 2.6972E-74 | 5.55258473 | 0.782 | 0.229 | 5.1128E-70 |
|  | IGLC3 | 1.9024E-25 | 5.08625896 | 0.354 | 0.065 | 3.6062E-21 |
|  | IGHG1 | 8.2738E-38 | 4.75648189 | 0.443 | 0.074 | 1.5684E-33 |
|  | IGLC2 | 8.4419E-42 | 4.67728928 | 0.483 | 0.071 | 1.6002E-37 |
|  | IGHG4 | 5.2891E-35 | 3.64364211 | 0.361 | 0.024 | 1.0026E-30 |
|  | IGHG3 | 3.6208E-25 | 3.36232375 | 0.319 | 0.047 | 6.8636E-21 |
|  | IGHA1 | 9.5936E-16 | 3.24707783 | 0.293 | 0.082 | 1.8186E-11 |
|  | IGHG2 | 3.7373E-09 | 2.66754242 | 0.182 | 0.056 | 7.0844E-05 |
|  | BANK1 | 3.2173E-46 | 1.97209874 | 0.517 | 0.118 | 6.0988E-42 |
| nmsccs | CD74 | 6.6924E-51 | 1.83636947 | 0.5 | 0.229 | 1.2686E-46 |
| Immune - subcluster5 | CD83 | 1.5424E-78 | 1.82935524 | 0.74 | 0.162 | 2.9237E-74 |
|  | JCHAIN | 9.8022E-10 | 1.81224322 | 0.18 | 0.05 | 1.8581E-05 |
|  | BACH2 | 7.0135E-36 | 1.68899281 | 0.413 | 0.068 | 1.3295E-31 |
|  | CCL3 | 1.4453E-21 | 1.68372075 | 0.325 | 0.074 | 2.7398E-17 |
|  | SIPA1L1 | 1.0015E-83 | 1.59421381 | 0.745 | 0.156 | 1.8984E-79 |
|  | FCHSD2 | 5.1503E-76 | 1.5504018 | 0.771 | 0.235 | 9.763E-72 |
|  | PDE4D | 3.3417E-34 | 1.54939227 | 0.429 | 0.091 | 6.3345E-30 |
|  | CCL3L3 | 6.6018E-15 | 1.54481846 | 0.257 | 0.068 | 1.2514E-10 |
|  | AFF3 | 8.4533E-45 | 1.5247373 | 0.57 | 0.174 | 1.6024E-40 |
|  | HLA-DQA1 | 5.6345E-79 | 1.51836775 | 0.77 | 0.203 | 1.0681E-74 |
|  | NAMPT | 2.0969E-30 | 1.5142951 | 0.464 | 0.129 | 3.9749E-26 |
| nmsccs | HLA-DRA | 1.409E-64 | 1.51000515 | 0.433 | 0.13 | 2.6708E-60 |
| Immune - subcluster5 | NFKB1 | 8.5166E-54 | 1.4969718 | 0.543 | 0.074 | 1.6144E-49 |
| fibro | SAA1 | 5.23E-148 | 1.47113224 | 0.11 | 0.013 | 9.913E-144 |
| Immune - subcluster5 | ANKRD44 | 3.1376E-68 | 1.45879327 | 0.728 | 0.221 | 5.9477E-64 |

**Supplementary Table 2.** Fraction of cells expressing *Toll-like receptors (TLR)* in the fibroblast cell cluster.

|  | Healthy pulp | Carious pulp |
| --- | --- | --- |
| TLR1 | 0.037986 | 0.052001 |
| TLR2 | 0.000968 | 0.004066 |
| TLR3 | 0.033143 | 0.070618 |
| TLR4 | 0.032067 | 0.065055 |
| TLR5 | 0.015711 | 0.035523 |
| TLR6 | 0.007425 | 0.015836 |
| TLR7 | 0.000323 | 0.001498 |
| TLR8 | 0.000323 | 0.000428 |
| TLR10 | 0.00043 | 0.00107 |

**Supplementary Table 3.**
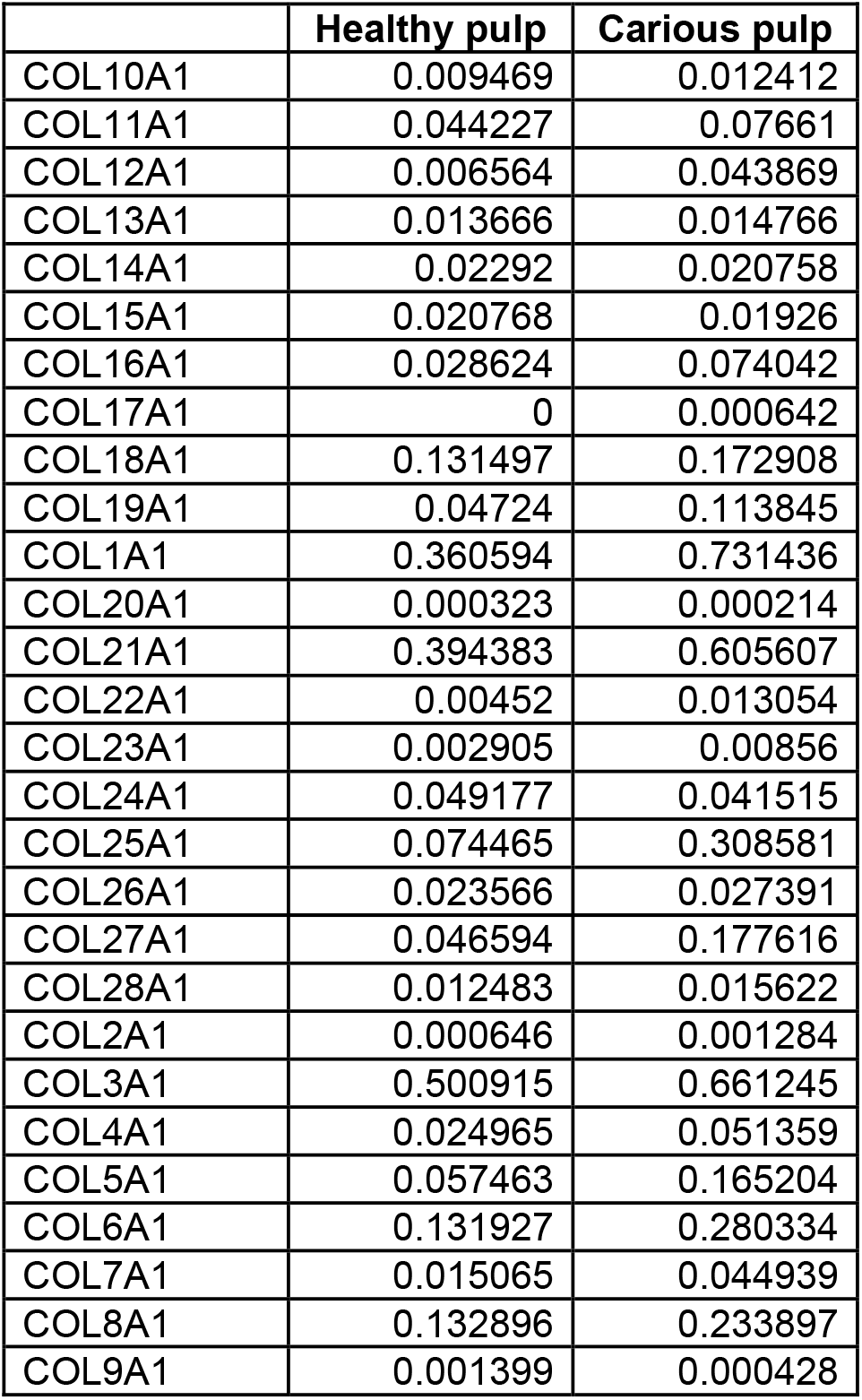
Fraction of cells expressing different Collagen types in the fibroblast cell cluster.

|  | **Healthy pulp** | **Carious pulp** |
| --- | --- | --- |
| COL10A1 | 0.009469 | 0.012412 |
| COL11A1 | 0.044227 | 0.07661 |
| COL12A1 | 0.006564 | 0.043869 |
| COL13A1 | 0.013666 | 0.014766 |
| COL14A1 | 0.02292 | 0.020758 |
| COL15A1 | 0.020768 | 0.01926 |
| COL16A1 | 0.028624 | 0.074042 |
| COL17A1 | 0 | 0.000642 |
| COL18A1 | 0.131497 | 0.172908 |
| COL19A1 | 0.04724 | 0.113845 |
| COL1A1 | 0.360594 | 0.731436 |
| COL20A1 | 0.000323 | 0.000214 |
| COL21A1 | 0.394383 | 0.605607 |
| COL22A1 | 0.00452 | 0.013054 |
| COL23A1 | 0.002905 | 0.00856 |
| COL24A1 | 0.049177 | 0.041515 |
| COL25A1 | 0.074465 | 0.308581 |
| COL26A1 | 0.023566 | 0.027391 |
| COL27A1 | 0.046594 | 0.177616 |
| COL28A1 | 0.012483 | 0.015622 |
| COL2A1 | 0.000646 | 0.001284 |
| COL3A1 | 0.500915 | 0.661245 |
| COL4A1 | 0.024965 | 0.051359 |
| COL5A1 | 0.057463 | 0.165204 |
| COL6A1 | 0.131927 | 0.280334 |
| COL7A1 | 0.015065 | 0.044939 |
| COL8A1 | 0.132896 | 0.233897 |
| COL9A1 | 0.001399 | 0.000428 |

